# The effect of autophagy and mitochondrial fission on Harderian gland is greater than apoptosis in male hamsters during different photoperiods

**DOI:** 10.1101/2020.05.26.116889

**Authors:** Zhe Wang, Jin-Hui Xu, Jun-Jie Mou, Xiang-Yu Zhao, Xiao-Cui Geng, Ming Wu, Hui-Liang Xue, Lei-Chen, Lai-Xiang Xu

## Abstract

Photoperiod is an important factor of mammalian seasonal rhythm. We studied the morphological differences in HG which is a vital photosensitive organ of male striped dwarf hamsters (*Cricetulus barabensis*), under different photoperiods (short photoperiod, SP; moderate photoperiod, MP; long photoperiod, LP), and further investigated the molecular mechanisms related to these morphological differences. Results showed that body weight, carcass weight, and HG weight were lower in SP and LP. Protein expression of bax/bcl2 and Cytochrome C showed no significant differences, indicating that the level of apoptosis remained stable. Protein aggregation of LC3 and protein expression of LC3II/LC3I were higher in SP than. Furthermore, comparison of changes in the HG ultrastructure demonstrated autolysosome formation in the LP, which suggesting the lowest autophagy level in MP. Protein expression levels of ATP synthase and mitochondrial fission factor were highest in the MP, whereas citrate synthase, dynamin-related protein1, and fission1 remained unchanged in three groups. In summary, the significant up-regulation of autophagy under short and long photoperiod may be the main factor leading to the loss of HG weight and reduced mitochondrial energy supply.

## Introduction

Seasonal rhythm is an adaptive behavior of animals in temperate regions to seasonal changes and includes changes in development, reproduction, hair growth, and energy metabolism (1,2). The HG, also known as the glandulae lacrimales accessoriae, covers the posterior part of two eyeballs and widely exists in mammals, birds, and reptiles (3,4); the HG weight of jungle bush quail (*Perdicula asiatica*) reached the highest in May, which showed a significant seasonal variation rhythm (5). In addition, seasonal reproductive behavior in animals is known to be impacted by changes in temperature, humidity, food resources, and photoperiod (i.e., length of sunshine), with photoperiod being one of the most important factors (6). However, whether seasonal variation in the HG is related to photoperiod remains to be clarified.

The balance between apoptosis and autophagy is one of important mechanism for tissue weight maintenance (7). Study on rats showed that the long-term light note leads to the increase of the level of apoptosis in HG (8,9). Bax is one of the most important apoptotic molecules in mammals and is activated under high mitochondrial depolarization for translocation and insertion into the outer membrane of mitochondria via bax/bax-homo-oligomerization (10). This is followed rapidly by the formation and opening of a mitochondrial permeability transition pore (mPTP), through which cytochrome C (Cyto C), a mitochondrion-residing apoptogenic factor, is released into the cytosol, eventually leading to the cleavage of nuclear DNA and cell apoptosis (11,12). At present, it is believed that DNA fragmentation detected by TUNEL staining is one of the most important indicators of increased apoptotic level (12). Furthermore, bcl2 inhibits apoptosis via suppression of bax/bax-homo-oligomerization (13,14). Research has shown that high-intensity light stimulation or high-dose MT injection can lead to increased cell necrosis in the HG of female Syrian hamsters and male rats (15). As MT is usually positively correlated with the time of entry into darkness (3,16,17), short daylight exposure may increase the level of apoptosis in the HG.Quantitative analysis of apoptosis in the HG can help clarify the mechanisms related to the effects of photoperiodic changes on the morphology and function of the HG.

Autophagy is the phagocytosis of cytoplasmic proteins or organelles and their entrapment and degradation in vesicles (18,19). As a key protein for autophagic lysosome formation, microtubule-associated protein 1 light chain (LC3-I) binds to the phosphatidylethanolamine (PE) complex to form LC3 II (20,21), which is a marker protein of intracellular macrophages as well as changes in autophagy (21,22). First discovered in 2013 (23), P62 is a transporter of degradable substances to autophagic lysosomes and is negatively related to autophagy levels in tissues (24). Quantitative analysis of LC3 and P62 proteins can indicate changes in autophagy level in the HG under different photoperiods. Melatonin can inhibit autophagy in the HG of female Syrian hamsters (25–27). To date, however, no research on the effects of photoperiod has been conducted in this field.

Changes in apoptotic and autophagic levels often involve mitochondrial function. Citrate synthase (CS) is a limiting enzyme of the tricarboxylic acid cycle (28,29) and adenosine triphosphate (ATP) synthase is a rate-limiting enzyme of the ATP synthesis pathway (30). Thus, studies on CS and ATP can partly measure changes in mitochondrial function and energy supply of the HG during different photoperiods. Changes in mitochondrial function may involve mitochondrial fission. Dynamin-related protein 1 (DRP1) is a GTP-hydrolyzing mechanoenzyme that catalyzes mitochondrial fission in the cell, which drives division via GTP-dependent constriction (31,32). The DRP1 receptor mitochondrial fission factor (Mff) is a major regulator of mitochondrial fission, with its overexpression resulting in increased fission (33). In contrast, DRP1 receptor fission 1 (FIS1) appears to recruit inactive forms of DRP1, and its overexpression inhibits mitochondrial fission (34,35). Therefore, research on these three factors could highlight mitochondrial fission ability. However, there is a current lack of research on mitochondrial fission and the function of HG during different photoperiods.

Based on the above, the effects of photoperiod on the HG may be related to autophagy, apoptosis, and mitochondrial function. Current photoperiod studies in hamsters have mainly focused on changes in HG morphology (36–38). However, the mechanisms involved in the morphological changes in the HG induced by different photoperiods, such as autophagy and apoptosis in small mammals during different photoperiods, remain poorly studied.The striped dwarf hamster (*Cricetulus barabensis*) is a small non-hibernating mammal widely distributed in the north temperate zone of Asia. In spring (March to April) and autumn (August to September), this hamster shows peak reproductive activities, while no reproductive activity occurs during the winter (December to January) (39,40). Our previous research have shown significant seasonal changes in gene expression like *kiss1*and *gpr54* in the hypothalamus, regulation of immune function, and energy metabolism in these animal (41). Thus, research on photoperiodic changes in this species of hamster could provide further insights into seasonal rhythm changes in non-hibernating mammals.

Here, we studied the morphological changes, as well as the related mechanisms, in the HG under different photoperiods. We hypothesized that photoperiodic changes would affect the morphology of the HG and thus its function. We also hypothesized that changes in apoptotic and autophagic levels may be responsible for changes in the HG. To test these hypotheses, we studied the ultrastructural changes in the HG of hamsters under different daylight lengths. On this basis, the protein levels of apoptosis (bax, bcl2, and Cyto C) and autophagy (LC3 and P62)-related indicators were studied. We then quantified mitochondrial function (ATP synthase and CS) and fission level (DRP1, MFF, and FIS1).

## Methods

### Ethics Statement

All procedures followed the Laboratory Animal Guidelines for the Ethical Review of Animal Welfare (GB/T 35892-2018) and were approved by the Animal Care and Use Committee of Qufu Normal University (Permit Number: dwsc 2019010).

### Animals and treatments

Striped dwarf hamsters were prepared as described previously in our laboratory (40,41). Briefly, hamsters were captured from the cropland in Qufu region of Shandong Province, China (N35.78°E117.01°). This area belongs to the temperate continental monsoon climate, light, and temperature changes with the seasons obviously. The main vegetation are wheat, peanuts and corn.

Captured hamsters were acclimated in the animal feeding room and exposed to natural light for about 2 weeks. Hamsters were housed individually in cages (28 ×18 ×12 cm) at an ambient temperature of 22 ±2 °C and relative humidity 55% ±5%. Food (standard rat chow, Jinan Pengyue Experimental Animal Breeding Co., Ltd., China) and water was provided *ad libitum*.

Based on the body weight and degree of wear on upper molars, a total of 60 male adult hamsters (20–40 g) were randomly divided into three groups of 20 animals. Each group were placed into long photoperiod (16:8 h light ? dark cycle; light from 04:00 to 20:00, LP), moderate photoperiod (12:12 h light ? dark cycle; light from 06:00 to 18:00, MP) or short photoperiod (8:16 h light ? dark cycle; light from 08:00 to 16:00, SP).

The hamsters for photoperiodic processing were placed in the cabin of biodiverse small animal feeding systems (NK, LP-30LED-8CTAR, Osaka, Japan). Temperature of cabin was 22 ±2 °C, relative humidity was 55% ±5% and light intensity was 150 ±10 lx. Photoperiodic processing lasted 8 weeks.

### Sample preparation

At the end of the exposure, hamsters were sacrificed by CO2 asphyxiation. The HGs were removed, with lengths and weights then recorded. The left HGs were immersed in glutaraldehyde-paraformaldehyde for transmission electron microscopy (TEM). The right HGs in each group were frozen in liquid nitrogen and stored in a refrigerator at −80 °C for subsequent Western blotting and immunofluorescence histochemical analyses. All procedures were carried out in accordance with approved guidelines.

### Transmission electron microscopy (TEM)

The HGs were cut into blocks and immersed in 3% glutaraldehyde-paraformaldehyde. The blocks were then dehydrated with a graded series of ethanol and embedded in epoxy resin, with TEM then performed as described previously (42). A semithin section was applied to tissue samples, and after methylene blue staining (18), sections were adjusted under the microscope and sliced with an ultramicrotome (LKB-NOVA, USA). The ultrathin sections were double-stained with Reynolds’ lead citrate and ethanolic uranyl acetate and then examined via TEM (JEOL, JEM-100SX, Japan). Images were processed with NIH Image software (Image-Pro Plus 6.0).

### Fluorescence immunohistochemical analysis

Frozen 10-μm thick tissue cross-sections were cut from the mid-belly of the two lobes each sample at −20 °C with a cryostat (Leica, CM1950, Germany) and then stored at −80 °C for further staining. Randomly selected 10 sections of each lobe for the followup experiments. After air drying for 2 h, endogenous peroxidase activity was inhibited in tissue sections using 0.5% v/v H2O2/methanol for 20 min at room temperature (43). Then after 15 min of immersion in distilled water, the sections were stained in blocking solution (5% bovine serum albumin (BSA)) for 30 min at room temperature and then incubated with rabbit anti-LC3 (1:200, #ab48394, Abcam, Cambridge, UK) or rabbit anti-P62 (1:200, #18420, Proteintech, Wuhan, China) solution at 4 °C overnight. The sections were subsequently incubated with goat anti-rabbit Alexa Fluor 488 (1:200, #11034, Thermo Fisher Scientific, Rockford, IL, USA) at 37 °C for 2 h Images were visualized using a confocal laser scanning microscope (ZEISS, 880NLO, Germany) under krypton/argon laser illumination at 488 nm emitted light, with capture at an emitting fluorescence of 526 nm. Protein aggregations of LC3 and P62 were counted using a 100 μm × 100 μm area.

### Terminal deoxynucleotidyl transferase biotin-dUTP nick end labeling (TUNEL) staining

DNA fragmentation induced by apoptosis was determined by double-labeled fluorometric TUNEL detection as described previously (10). The frozen sections were permeabilized with 0.2% Triton X-100 in 0.1% sodium citrate at 4 °C for 2 min and then incubated with an anti-laminin rabbit polyclonal antibody (1:500, #BA1761, Boster, Wuhan, China) at 4 °C overnight. After washing with PBS for 30 min, the sections were incubated with fluorochrome-conjugated secondary AF647 antibodies (1:200, #21245, Thermo Fisher Scientific) at room temperature for 2 h. Subsequently, TUNEL (#MK1023, Boster) reaction mixture was added at the recommended 1:9 ratio, and the sections were incubated for 60 min at 37 °C in a humidified chamber in the dark, as per the manufacturer’s protocols. Finally, the sections were counterstained with DAPI (1:100, #D1306, Sigma-Aldrich, Saint Quentin Fallavier, France) at 37 °C for 30 min. Imaging was performed using a confocal laser scanning microscope with the same excitation and emission wavelengths as described above.

### Western blotting

Western blotting was conducted as described previously (44). Protein was extracted from HGs and solubilized in sample buffer (100 mM Tris pH 6.8, 5% 2-β-mercaptoethanol, 5% glycerol, 4% SDS, and bromophenol blue), with protein extracts subsequently fractionated by SDS-PAGE using Laemmli gels and transferred to PVDF membranes (0.45-μm pore size) using a Bio-Rad semi-dry transfer apparatus. The blotted membranes were blocked with 1% BSA in Tris-buffered saline (TBS; 150 mM NaCl, 50 mM Tris-HCl, pH 7.5) and incubated with rabbit anti-bax (1:1000, #50599, Proteintech), rabbit anti-bcl2 (1:1000, #3498, Cell Signaling Technology CST, Danvers, MA, USA), rabbit anti-Cyto C (1:1000, #11940, CST), rabbit anti-LC3 (1:1000), rabbit anti-P62 (1:1000) and rabbit rabbit anti-ATP synthase (1:1000, #14676, Proteintech), rabbit anti-citrate synthase (1:1000, #16131, Proteintech), rabbit anti-DRP1 (1:1000, #12957, Proteintech), rabbit anti-MFF (1:1000, #17090, Proteintech), rabbit anti-FIS1 (1:1000, #10956, Proteintech), and anti-β-actin (1:5000, #20536, Proteintech) in TBS containing 0.1% BSA at 4 °C overnight. The membranes were then incubated with IRDye 800 CW goat-anti rabbit secondary antibodies (1:5000, #31460, Thermo Fisher Scientific) for 90 min at room temperature and visualized with an Odyssey scanner (Bio-Rad, California, USA). Quantification analysis of the blots was performed using NIH Image J software.

### Statistical analyses

The normality of data and homogeneity of variance are tested by Shapiro-Wilk and Levene respectively. Single factor analysis of variance (one-way ANOVA) was used to compare the differences between groups. When the variance is homogeneous, the least significant difference (LSD) post-hoc test was used for multiple comparisons among groups. When the variance is not homogeneous, the Dunnett T3 method was used for comparison among groups. The differences are considered significant when *P* < 0.05. Data are expressed as means ±standard deviation (Mean ±SD). All statistical analyzes were conducted using SPSS 19.0.

## Results

### Changes in Harderian gland weight (HGW) and Harderian gland weight to carcass weight ratio (HGW/CW) in hamsters under different photoperiods

The HGW was significantly lower in SP (5%, *P* < 0.05) and LP (5%, *P* < 0.05) group than MP group, while the HGW/CW ratio demonstrated no significant differences among the three groups (Fig. 1).

**Fig. 1.**
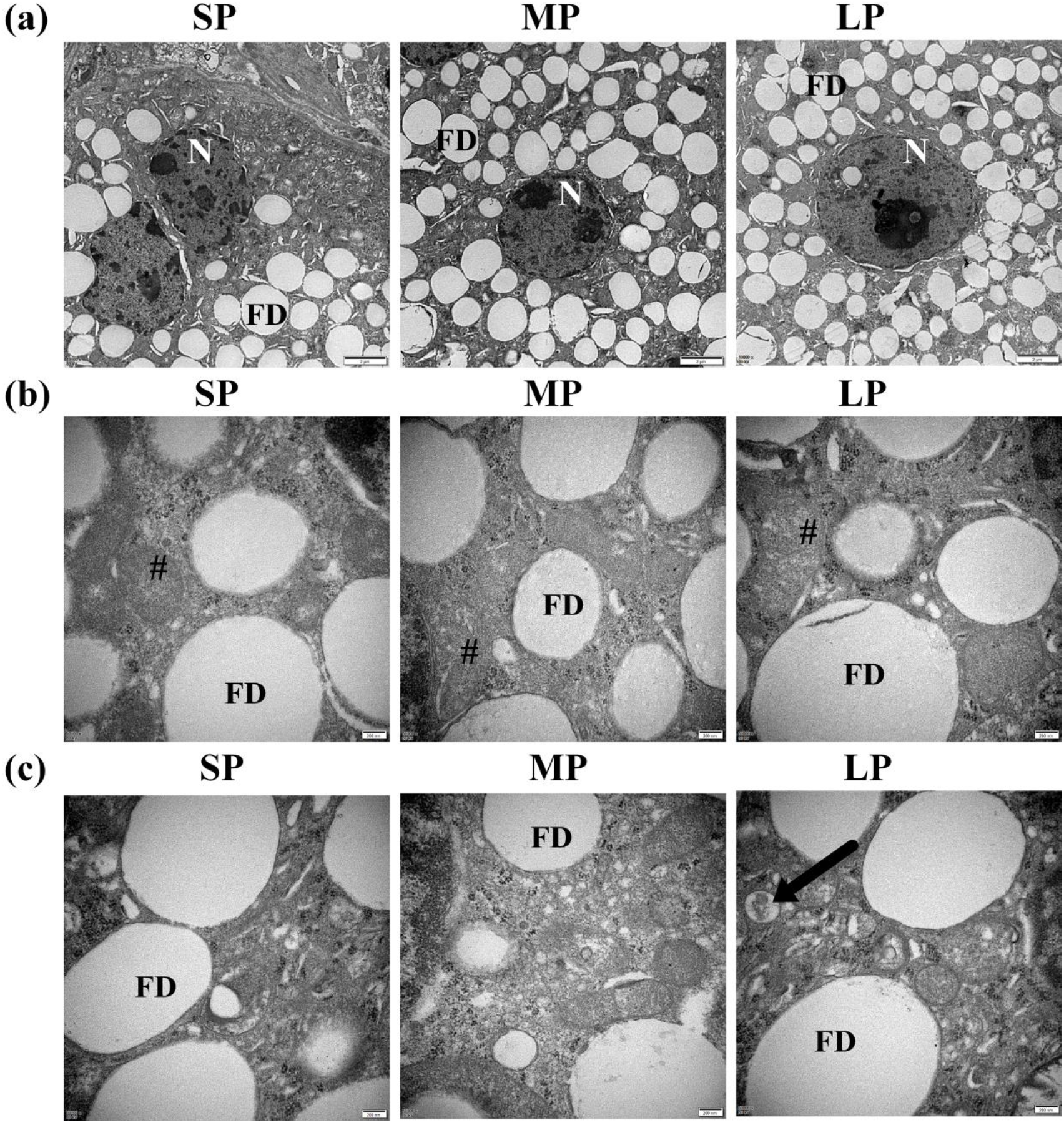
Ultrastructure of HG in hamsters from three photoperiodic groups. (a) Nucleus ultrastructure of HG in hamsters from three photoperiodic groups. There were no significant differents in nuclear (N) morphology among three photoperiodic groups. Large number of fat droplets (FD) were observed in secretory cells of HG. Scale bar = 2 μm. (b) Cristae of mitochondria of HG in hamsters from three photoperiodic groups. There were no significant differents in mitochondrial (#) morphology among three groups. Scale bar = 0.2 μm. (C) Autophagolysosomes of HG in hamsters from three photoperiodic groups. Significant autophagolysosomal structures (see arrow) were observed in LP group. In other groups, autophagolysosomal structures were hardly observed. Scale bar = 0.2 μm. SP, short photoperiod; MP, moderate photoperiod; LP, long photoperiod.

### Ultrastructural changes in HG nuclei, mitochondria, and autophagolysosomes

A large number of secretory cells were observed in the HGs of the three different photoperiod groups, including a large number of round- or elliptical-shaped fat droplets. The plasma membrane of the secretory cells was clearly visible. The mitochondria in the HG were irregularly oval, the cristae were clearly visible, and the membrane was complete. There was no significant difference in nuclear and mitochondrial morphology among the three groups. Typical autophagolysosomal structures were observed in the LD group, showing a clear membrane structure on the outside and wrapped contents in the middle. In other groups, however, it was difficult to observe typical autophagolysosomal structures (Fig. 2).

**Fig. 2.**
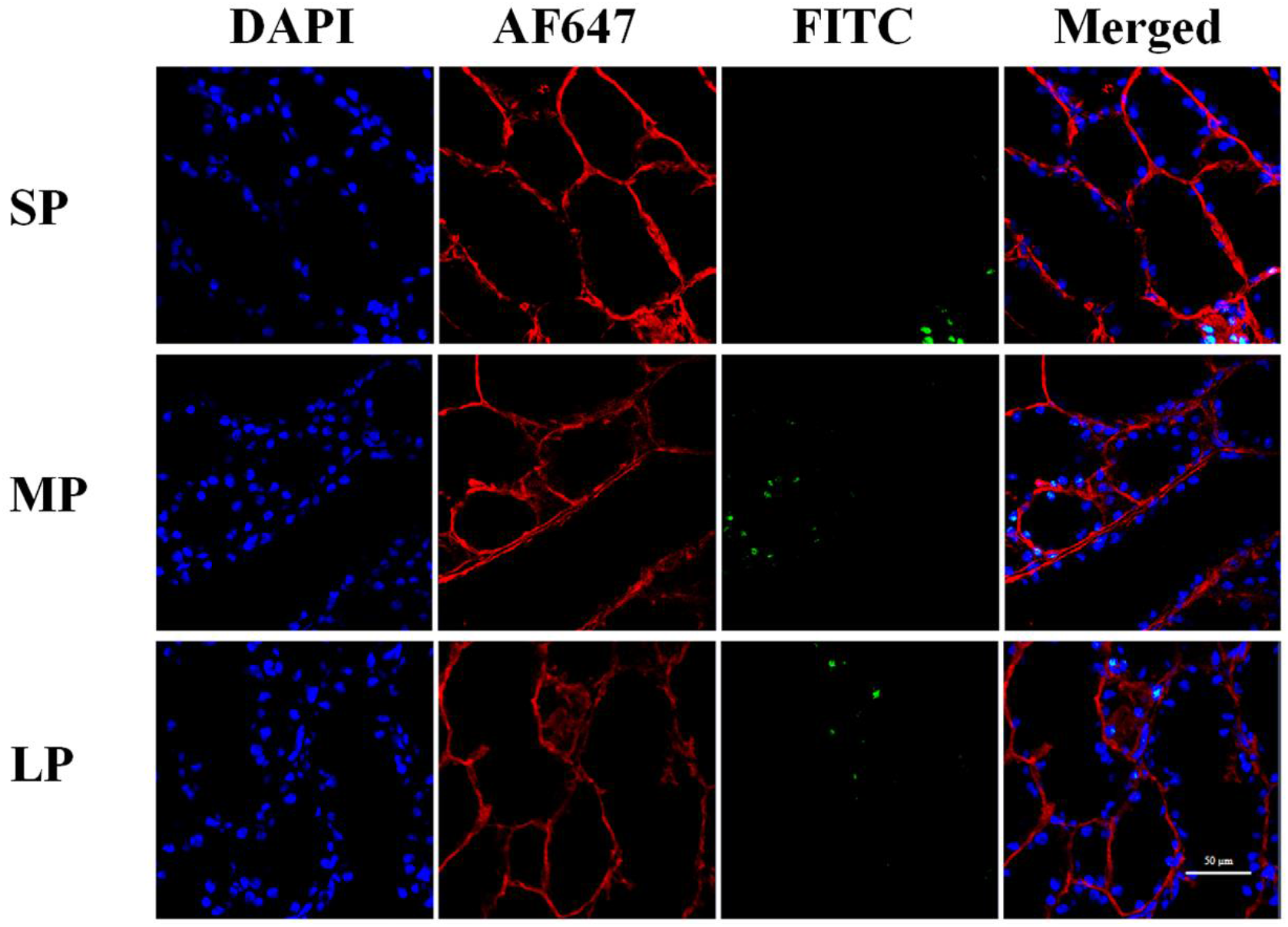
Fluorescent terminal deoxynucleotidyl transferase biotin-dUTP nick end labeling (TUNEL) staining of HG in hamsters in three photoperiodic groups. Immunofluorescence histochemistry showing cell apoptosis, cell boundaries, and nuclei. Blue represents 4’6’-diamidino-2-phenylindole (DAPI)-stained nucleus, red represents Alexa Fluor 647-stained laminin of interstitial tissue, green represents TUNEL by FITC. Scale bar = 50 μm. SP, short photoperiod; MP, moderate photoperiod; LP, long photoperiod.

### DNA fragmentation

TUNEL staining provided direct evidence of apoptosis. In three photoperiodic groups, random HG sections showed that DNA fragmentation (represented by green fluorescence) were observed (Fig. 3).

**Fig. 3.**
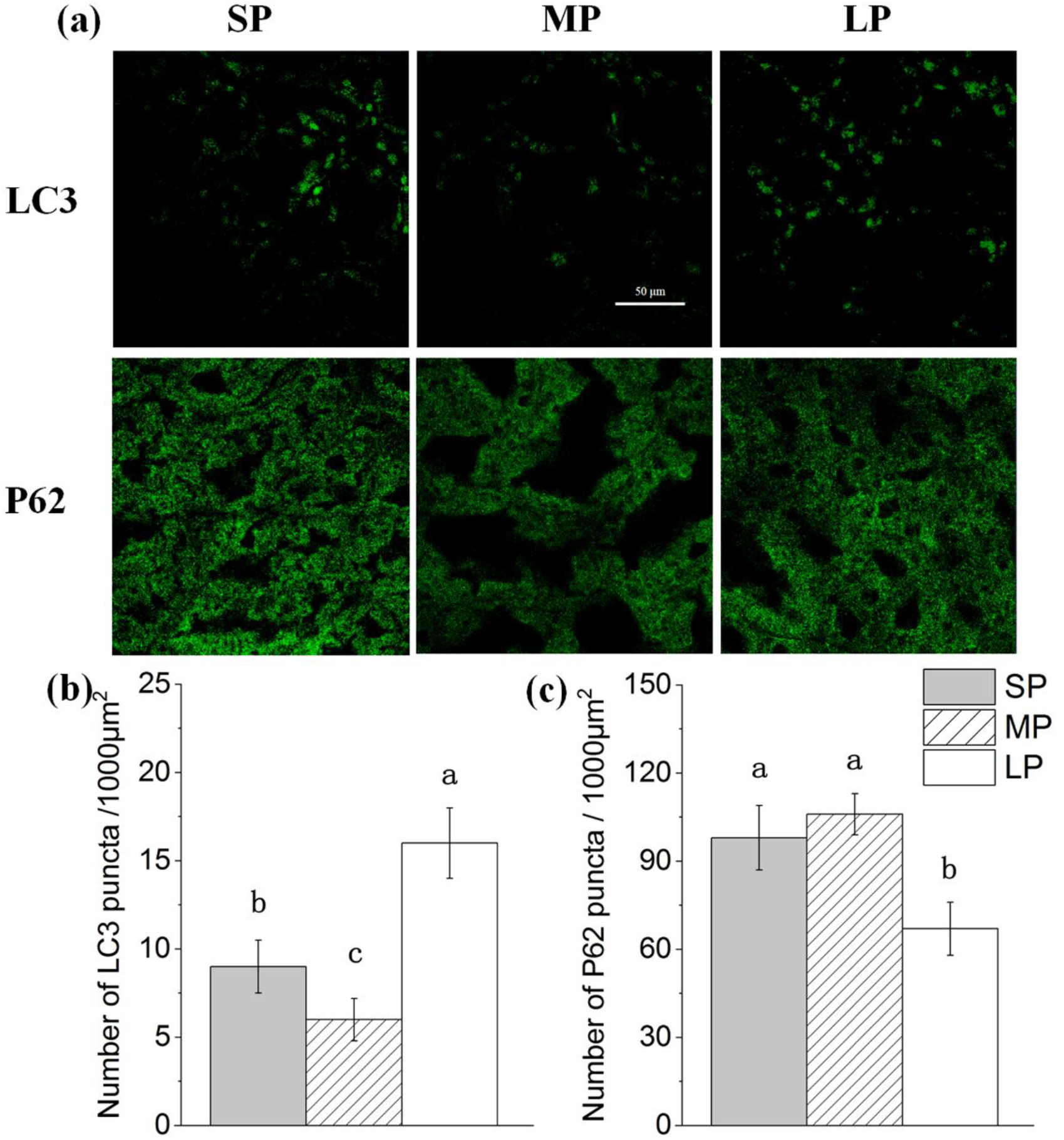
Quantification of LC3 and P62 puncta of HG in hamsters in three photoperiodic groups. (a) Immunofluorescence histochemistry showing LC3 and P62 puncta. green represents AF488-stained LC3 or P62, respectively. Scale bar = 50 μm. (b) Quantification of LC3 puncta. (c) Quantification of P62 puncta. Six figures were analyzed in each sample; ten samples were analyzed in each group. Values are means ±SD. SP, short photoperiod; MP, moderate photoperiod; LP, long photoperiod. Different letters identify statistically significant difference (*P* < 0.05).

### Changes in LC3 and P62 puncta in HG under different photoperiods

The number of cytoplasmic LC3 puncta per 1 000 μm^2^ is indicative of LC3I to LC3II conversion. Representative figures of LC3 immunofluorescence staining are shown in Fig. 4a, which significantly higher 77% and 172% (*P* < 0.05) in the SP and LP group, respectively, than that in the MP group (Fig. 4b).

**Fig. 4.**
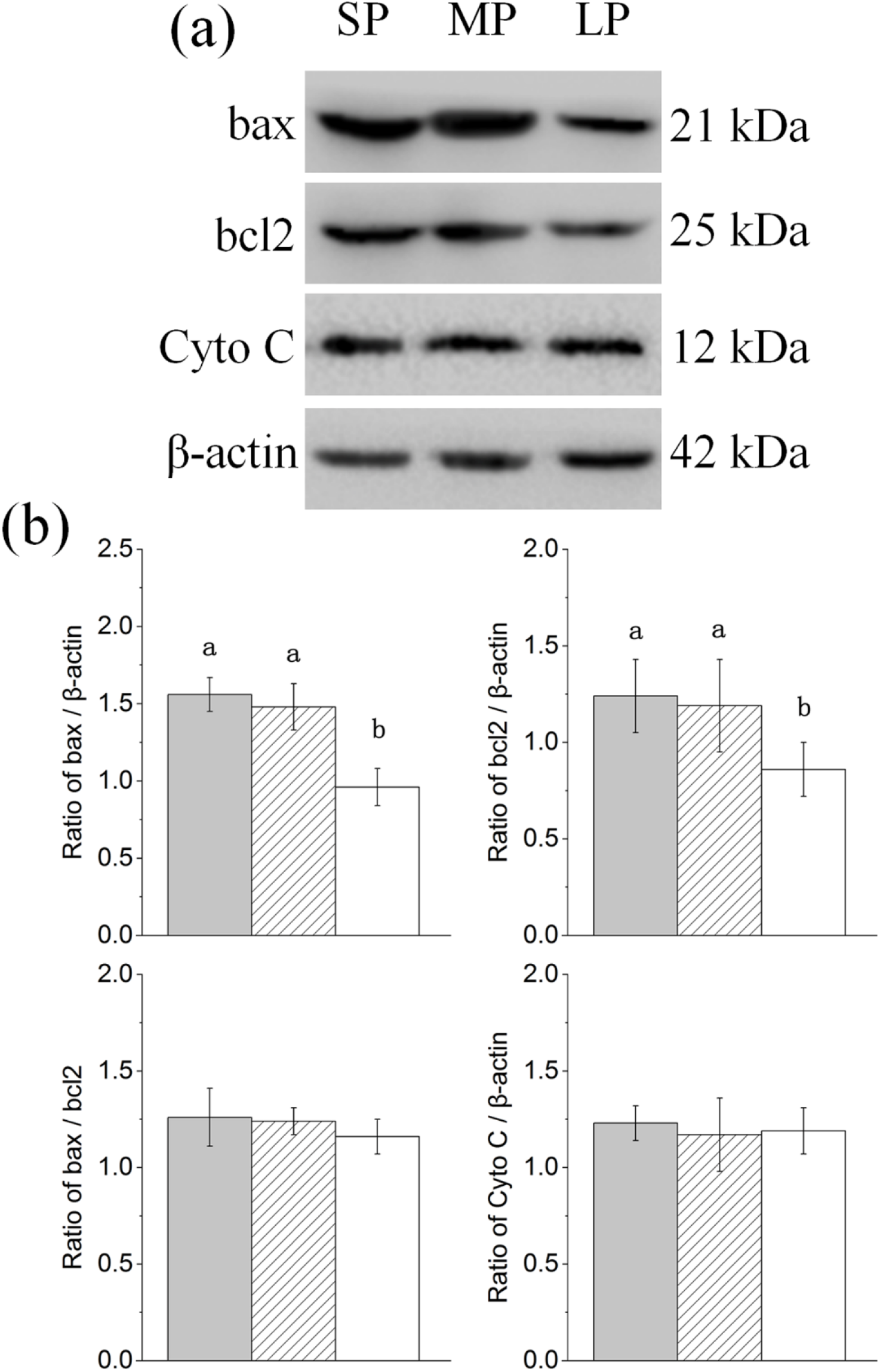
Changes in protein levels of apoptosis related factors in HG of hamsters in three different photoperiodic groups. (a) Representative immunoblots of bax, bcl2, Cyto C, and β-actin in three different photoperiodic groups. (b) Ratio of bax, bcl2, Cyto C to β-actin and ratio of bax to bcl2 in HG of hamsters in three different photoperiodic groups. Values are means ±SD. n=10. SP, short photoperiod; MP, moderate photoperiod; LP, long photoperiod. Different letters identify statistically significant difference (*P* < 0.05).

Representative figures of P62 immunofluorescence staining are shown in Fig. 4a The number of P62 puncta was lowest in the LP group, with 43% (*P* < 0.05) less P62 puncta in the MP group (Fig. 4c).

### Relative protein expression of apoptosis related factors

The contents of bax, bcl2, and Cyto C were detected by Western blot analysis, as shown in Figure 5a. The ratio of bax/bcl2 and the protein expression of Cyto C all showed no significant difference between three groups (Fig. 5b).

**Fig. 5.**
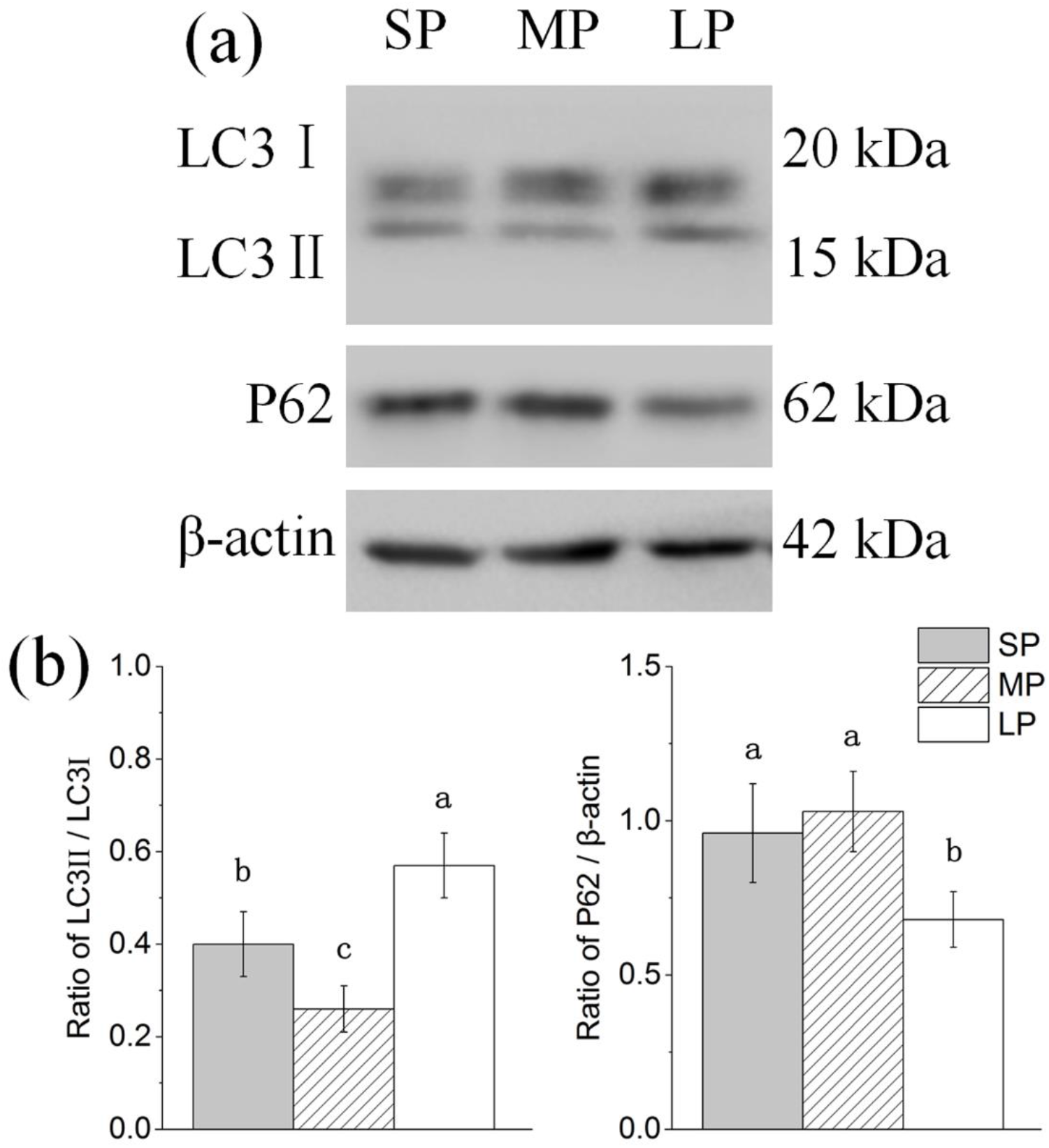
Changes in protein levels of autophagy related factors in HG of hamsters in three different photoperiodic groups. (a) Representative immunoblots of LC3, P62, and β-actin in three different photoperiodic groups. (b) Ratio of LC3, P62 to β-actin in HG of hamsters in three different photoperiodic groups. Values are means ±SD. n=10. SP, short photoperiod; MP, moderate photoperiod; LP, long photoperiod. Different letters identify statistically significant difference (*P* < 0.05).

### Relative protein expression of autophagy related factors

The contents of LC3 and P62 were detected by Western blot analysis, as shown in Figure 6a. The LC3II/LC3I level showed higher in SP (43%, *P* < 0.05) and LP groups (113%, *P* < 0.05) than that in MP group. Meanwhile, the protein expression of P62 showed LP group was lower (*P* < 0.05) than SP and MP groups (Fig. 6b).

**Fig. 6.**
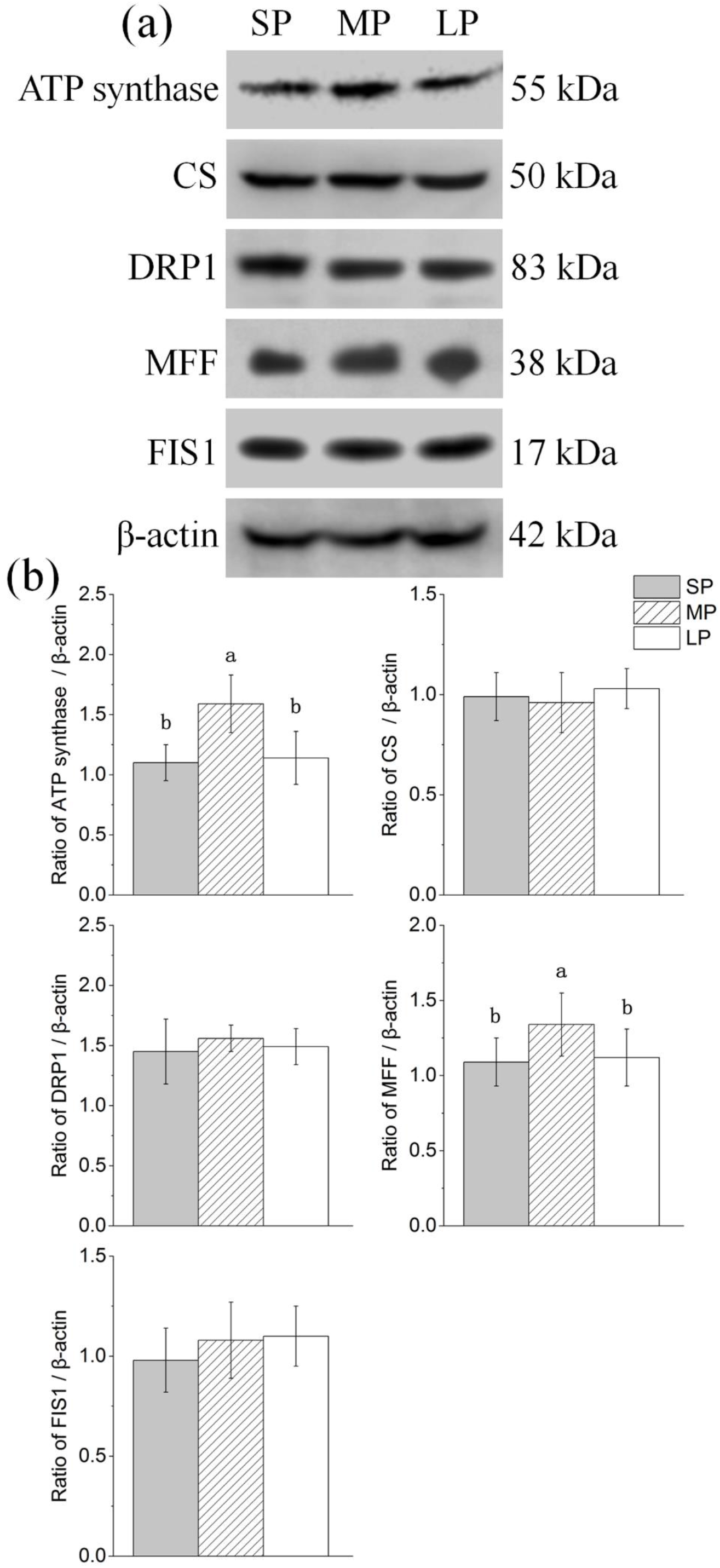
Changes in protein levels of mitochondrial related factors in HG of hamsters in three different photoperiodic groups. (a) Representative immunoblots of ATP synthase, CS, DRP1, MFF, FIS1, and β-actin in three different photoperiodic groups. (b) Ratio of ATP synthase, CS, DRP1, MFF, FIS1 to β-actin in HG of hamsters in three different photoperiodic groups. Values are means ±SD. n = 10. SP, short photoperiod; MP, moderate photoperiod; LP, long photoperiod. Different letters identify statistically significant difference (*P* < 0.05).

### Relative protein expression of mitochondrial related factors

The contents of ATP synthase, CS, DRP1, MFF, and FIS1 were detected by Western blot analysis, as shown in Figure 7a. CS, DRP1, and FIS1 protein expression showed no significant differences among the three groups. ATP synthase and MFF protein expression in the MD group was significantly increased (*P* < 0.05) compared with that in the other two groups (Fig. 7b).

**Fig. 7.**
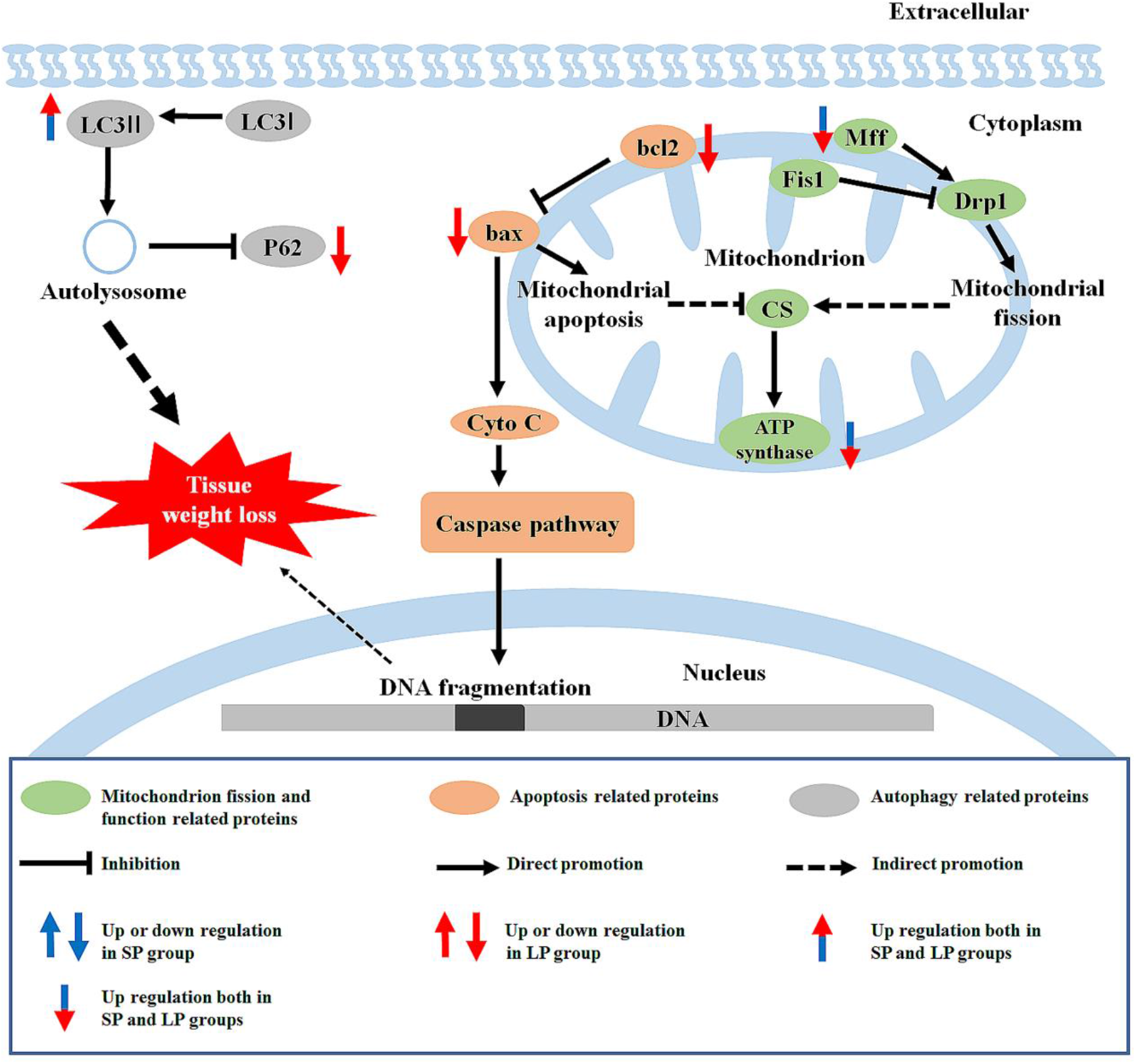
Graphical summary of study. LC3, microtubule-associated protein 1 light chain; P62, sequestosome 1; Cyto C, cytochrome C; FIS1, fission 1; MFF, mitochondrial fission factor; DRP1, dynamin- related protein 1; ATP synthase, adenosine triphosphate synthase; CS, citrate synthase; SP, short photoperiod; LP, long photoperiod.

## Discussion

Our results showed that, compared with the moderate photoperiod control group, the HG weight of hamsters was significantly reduced under short and long photoperiods. The protein expression levels of bax/bcl2 and Cyto C also showed no significant differences among groups. In contrast, the protein expression of LC3II/LC3I was higher in the short and long photoperiod groups compared with the moderate photoperiod control. Furthermore, protein expression of ATP synthase and MFF was highest in the moderate photoperiod control group.

We found that after 10 weeks of different light treatment, HG weight in the short and long photoperiod groups was lower than that in the moderate photoperiod group.

However, the HG weight to carcass weight ratio in the short and long photoperiod groups was slight change, suggesting that the decrease in HG weight may be consistent with a change in animal carcass weight.

To explore the mechanism of the above phenomenon, we studied the apoptosis level in HG of hamsters under different photoperiods and found that DNA fragmentation occurred in all three groups and ultrastructure analysis also showed that there was no significant nuclear change in the secretory cells of the HG during three groups. The bax/bcl2 ratio is often used to measure the degree of cell apoptosis and Cyto C is the key signal of apoptosis. (13,14). Our study showed that although bax decreased significantly in the long photoperiod group, the ratio of bax to bcl2 and the protein expression of Cyto C remained stable, indicating that the level of apoptosis might be stable among the three groups. High-intensity light stimulation or high-dose MT injection can lead to increased cell necrosis in the HG of female Syrian hamsters and male rats (15). Since melatonin is usually positively correlated with the time to enter darkness (3,16,17), this may mean that short photoperiod exposure may increase the apoptosis level in HG. However, our research did not find this. On the one hand, it may be that exogenous injection of MT is not the same as simple photoperiod treatment. On the other hand, Syrian hamsters hibernate in winter (45), whereas striped dwarf hamsters display daily torpor (46). Therefore, the effects on these two hamsters differ under the short photoperiod of winter.

Interestingly, we found that the protein expression of LC3II/LC3I was higher in the short and long photoperiod groups than that in the moderate photoperiod control. As LC3II is a key protein of autophagolysosome membrane formation (20,21), this result indicates that the level of autophagy may be higher in these two groups. In addition, the number of cytoplasmic LC3 puncta, which is indicative of LC3I to LC3II conversion, showed the same trend as LC3II/LC3I protein expression, which also suggests an increase in autophagy level. P62 is an autophagic transport protein, the accumulation of which indicates a decrease in autophagic efficiency (24). Here, immunofluorescence histochemistry showed that P62 protein aggregates and expression levels were lower in the long photoperiod group than in the other two groups, indicating that the level of autophagy might be highest under long photoperiod conditions. This is consistent with the ultrastructural results, showing the occurrence of autolysosomes. As the balance between apoptosis and autophagy is an important mechanism for tissue weight maintenance (7), the higher autophagy level under the short and long photoperiods compared to the moderate photoperiod may be the main reason for the lower HG weight in hamsters.

We also found that ATP synthase protein expression levels were lower following short and long photoperiod treatment, consistent with the change in mitochondrial fission level. CS is a rate-limiting enzyme in the tricarboxylic acid cycle and represents the ability of mitochondria to undertake aerobic oxidation (28,29). ATP synthase is the last step in ATP production by mitochondria, representing the ability of mitochondria to supply energy (30). In this study, the protein expression levels of ATP synthase in the HG were lower under short and long photoperiod conditions, whereas CS was maintained, indicating that the mitochondrial energy supply function was weakened slightly, and mitochondrial aerobic capacity remained unchanged. As ultrastructural analysis showed that mitochondria remained relatively intact in the three groups, we speculated that mitochondrial function may not have been weakened significantly. Drp1 is a key factor related to the promotion of mitochondrial fission, with MFF and FIS1 found to up- and down-regulate DRP1 activity, respectively (33,35). In the short and long photoperiod groups, MFF protein expression decreased, whereas that of DRP1 and FIS1 remained unchanged. Considering that MFF is the up-regulatory factor of mitochondrial fission, rather than the most important factor, these results indicate that the mitochondrial fission level may have decreased slightly, which could be one of the reasons for the slight decrease in mitochondrial energy supply.

In summary, this study extends novel findings on the effects of photoperiod on morphological and functional changes in the HG and related mechanisms under different photoperiods (Fig. 7). As there were no significant changes in the level of apoptosis in the HG under the different photoperiods, the significant up-regulation in autophagy level under long and short photoperiod conditions may be the main factor leading to tissue weight loss. Mitochondrial function weakened slightly under short and long photoperiod treatment, which may be caused by the maintenance of apoptosis and down-regulation of mitochondrial fission. Photoperiod treatment in the non-breeding season (i.e., short and long photoperiods) led to different levels of degeneration in the morphology and function of the HG in hamsters, with the possible mechanism involving autophagy and mitochondrial fission.

**Table 1.** Effects of photoperiod on carcass weight (CW), Harderian gland wet weight (HGWW), and ratio of HGWW/CW in hamsters after 10 weeks

| Group | SP | MP | LP |
| --- | --- | --- | --- |
| CW after photoperiod (g) | 15.91±1.26 | 17.06±2.51 | 15.89±0.84 |
| HGWW after photoperiod (mg) | 22.5±1.83<sup>b</sup> | 25.1±1.57<sup>a</sup> | 22.9 ±1.5<sup>b</sup> |
| HGWW/CW after photoperiod (mg/g) | 1.42 ±0.12 | 1.47 ±0.07 | 1.44 ±0.07 |

Values are means ± SD. n = 10. SP, short photoperiod; MP, moderate photoperiod; LP, long photoperiod. Different letters identify statistically significant difference ( $P < 0.05$ ).

## Availability of data and materials

The datasets used and/or analyzed during the current study are available from the corresponding author on reasonable request. Partial original images are included in the supplementary documents.

## Acknowledgments

Not applicable.

## Author Contributions

ZW, JX, and LX conceived and designed research; ZW, JX, JM, and XZ performed experiments; ZW analyzed data; ZW interpreted experimental results; ZW and JM prepared figures; ZW and JX drafted manuscript; MW and HX provided experimental guidance and suggestions for revision; JX, ZW, and LX edited manuscript and approved final version of manuscript.

## Consent for publication

Not applicable.

## References

1. Nakayama, T., and Yoshimura, T. (2018) Seasonal Rhythms: The Role of Thyrotropin and Thyroid Hormones. Thyroid: official journal of the American Thyroid Association. 28, 4–10

2. Masson-Pevet, M., Naimi, F., Canguilhem, B., Saboureau, M., Bonn, D., and Pevet, P. (1994) Are the annual reproductive and body weight rhythms in the male European hamster (Cricetus cricetus) dependent upon a photoperiodically entrained circannual clock? Journal of pineal research 17, 151–163

3. Buzzell, G. R. (1996) The Harderian gland: perspectives. Microscopy research and technique 34, 2–5

4. Sakai, T. (1989) Major ocular glands (harderian gland and lacrimal gland) of the musk shrew (Suncus murinus) with a review on the comparative anatomy and histology of the mammalian lacrimal glands. Journal of morphology 201, 39–57

5. Dubey, S., and Haldar, C. (1997) Environmental factors and annual harderian-pineal-gonadal interrelationship in Indian jungle bush quail, Perdicula asiatica. General and comparative endocrinology 106, 17–22

6. Borniger, J. C., and Nelson, R. J. (2017) Photoperiodic regulation of behavior: Peromyscus as a model system. Seminars in cell & developmental biology 61, 82–91

7. Gonzalez, C. R., Isla, M. L. M., and Vitullo, A. D. (2018) The balance between apoptosis and autophagy regulates testis regression and recrudescence in the seasonal-breeding South American plains vizcacha, Lagostomus maximus. Plos One 13

8. Monteforte, R., Santillo, A., Lanni, A., D’Aniello, S., and Baccari, G. C. (2008) Morphological and biochemical changes in the Harderian gland of hypothyroid rats. J Exp Biol 211, 606–612

9. Kurisu, K., Sawamoto, O., Watanabe, H., and Ito, A. (1996) Sequential changes in the harderian gland of rats exposed to high intensity light. Laboratory animal science 46, 71–76

10. Fu, W. W., Hu, H. X., Dang, K., Chang, H., Du, B., Wu, X., and Gao, Y. F. (2016) Remarkable preservation of Ca2+ homeostasis and inhibition of apoptosis contribute to anti-muscle atrophy effect in hibernating Daurian ground squirrels. Sci Rep-Uk 6, 13

11. Zha, H., Aime-Sempe, C., Sato, T., and Reed, J. C. (1996) Proapoptotic protein Bax heterodimerizes with Bcl-2 and homodimerizes with Bax via a novel domain (BH3) distinct from BH1 and BH2. The Journal of biological chemistry 271, 7440–7444

12. Smith, H. K., Maxwell, L., Martyn, J. A., and Bass, J. J. (2000) Nuclear DNA fragmentation and morphological alterations in adult rabbit skeletal muscle after short-term immobilization. Cell and tissue research 302, 235–241

13. Antonsson, B., Conti, F., Ciavatta, A., Montessuit, S., Lewis, S., Martinou, I., Bernasconi, L., Bernard, A., Mermod, J. J., Mazzei, G., Maundrell, K., Gambale, F., Sadoul, R., and Martinou, J. C. (1997) Inhibition of Bax channel-forming activity by Bcl-2. Science 277, 370–372

14. Korsmeyer, S. J., Shutter, J. R., Veis, D. J., Merry, D. E., and Oltvai, Z. N. (1993) Bcl-2/Bax: a rheostat that regulates an anti-oxidant pathway and cell death. Seminars in cancer biology 4, 327–332

15. Vega-Naredo, I., Caballero, B., Sierra, V., Garcia-Macia, M., de Gonzalo-Calvo, D., Oliveira, P. J., Rodriguez-Colunga, M. J., and Coto-Montes, A. (2012) Melatonin modulates autophagy through a redox-mediated action in female Syrian hamster Harderian gland controlling cell types and gland activity. J. Pineal Res. 52, 80–92

16. Tsutsui, K., and Ubuka, T. (2018) Photoperiodism in Mammalian Reproduction,

17. Payne, A. P. (1994) The harderian gland: a tercentennial review. Journal of anatomy 185 (Pt 1), 1–49

18. Biazik, J., Vihinen, H., Anwar, T., Jokitalo, E., and Eskelinen, E. L. (2015) The versatile electron microscope: an ultrastructural overview of autophagy. Methods (San Diego, Calif.) 75, 44–53

19. Dong, S., Zhao, S., Wang, Y., Pang, T., and Ru, Y. (2015) [Analysis of blood cell autophagy distribution in hematologic diseases by transmission electron microscope]. Zhonghua xue ye xue za zhi = Zhonghua xueyexue zazhi 36, 144–147

20. Lee, Y. K., and Lee, J. A. (2016) Role of the mammalian ATG8/LC3 family in autophagy: differential and compensatory roles in the spatiotemporal regulation of autophagy. BMB reports 49, 424–430

21. Schaaf, M. B., Keulers, T. G., Vooijs, M. A., and Rouschop, K. M. (2016) LC3/GABARAP family proteins: autophagy-(un)related functions. FASEB journal: official publication of the Federation of American Societies for Experimental Biology 30, 3961–3978

22. Mizushima, N., and Yoshimori, T. (2007) How to interpret LC3 immunoblotting. Autophagy 3, 542–545

23. Zhang, Y. B., Gong, J. L., Xing, T. Y., Zheng, S. P., and Ding, W. (2013) Autophagy protein p62/SQSTM1 is involved in HAMLET-induced cell death by modulating apotosis in U87MG cells. Cell Death Dis 4

24. Lamark, T., Svenning, S., and Johansen, T. (2017) Regulation of selective autophagy: the p62/SQSTM1 paradigm. Essays in biochemistry 61, 609–624

25. Kongsuphol, P., Mukda, S., Nopparat, C., Villarroel, A., and Govitrapong, P. (2009) Melatonin attenuates methamphetamine-induced deactivation of the mammalian target of rapamycin signaling to induce autophagy in SK-N-SH cells. J Pineal Res 46, 199–206

26. Nopparat, C., Porter, J. E., Ebadi, M., and Govitrapong, P. (2010) The mechanism for the neuroprotective effect of melatonin against methamphetamine-induced autophagy. J Pineal Res 49, 382–389

27. Coto-Montes, A., and Tomas-Zapico, C. (2006) Could melatonin unbalance the equilibrium between autophagy and invasive processes? Autophagy 2, 126–128

28. Wiegand, G., and Remington, S. J. (1986) Citrate synthase: structure, control, and mechanism. Annual review of biophysics and biophysical chemistry 15, 97–117

29. Remington, S. J. (1992) Structure and mechanism of citrate synthase. Current topics in cellular regulation 33, 209–229

30. Kramarova, T. V., Shabalina, I. G., Andersson, U., Westerberg, R., Carlberg, I., Houstek, J., Nedergaard, J., and Cannon, B. (2008) Mitochondrial ATP synthase levels in brown adipose tissue are governed by the c-Fo subunit P1 isoform. Faseb Journal 22, 55–63

31. Kraus, F., and Ryan, M. T. (2017) The constriction and scission machineries involved in mitochondrial fission. Journal of cell science 130, 2953–2960

32. Tilokani, L., Nagashima, S., Paupe, V., and Prudent, J. (2018) Mitochondrial dynamics: overview of molecular mechanisms. Essays in biochemistry 62, 341–360

33. Tieu, Q., and Nunnari, J. (2000) Mdv1p is a WD repeat protein that interacts with the dynamin-related GTPase, Dnm1p, to trigger mitochondrial division. The Journal of cell biology 151, 353–366

34. Liu, R., and Chan, D. C. (2015) The mitochondrial fission receptor Mff selectively recruits oligomerized Drp1. Molecular biology of the cell 26, 4466–4477

35. Yu, R., Jin, S. B., Lendahl, U., Nister, M., and Zhao, J. (2019) Human Fis1 regulates mitochondrial dynamics through inhibition of the fusion machinery. EMBO J 38

36. Vaughan, M. K., Menendez-Pelaez, A., Buzzell, G. R., Vaughan, G. M., Little, J. C., and Reiter, R. J. (1994) Circadian rhythms in reproductive and thyroid hormones in gonadally regressed male hamsters exposed to natural autumn photoperiod and temperature conditions. Neuroendocrinology 60, 96–104

37. Menendez-Pelaez, A., Lopez-Gonzalez, M. A., and Guerrero, J. M. (1993) Melatonin binding sites in the Harderian gland of Syrian hamsters: sexual differences and effect of castration. J. Pineal Res. 14, 34–38

38. Buzzell, G. R., Blank, J. L., Vaughan, M. K., and Reiter, R. J. (1995) Control of secretory lipid droplets in the harderian gland by testosterone and the photoperiod: comparison of two species of hamsters. General and comparative endocrinology 99, 230–238

39. Xu, L. X., Xue, H. L., Li, S. N., Xu, J. H., and Chen, L. (2017) Seasonal differential expression of KiSS-1/GPR54 in the striped hamsters (Cricetulus barabensis) among different tissues. Integr. Zool. 12, 260–268

40. Xue, H. L., Xu, J. H., Chen, L., and Xu, L. X. (2014) Genetic variation of the striped hamster (Cricetulus barabensis) and the impact of population density and environmental factors. Zool. Stud. 53, 8

41. Zhao, L., Zhong, M., Xue, H. L., Ding, J. S., Wang, S., Xu, J. H., Chen, L., and Xu, L. X. (2014) Effect of RFRP-3 on reproduction is sex- and developmental statusdependent in the striped hamster (Cricetulus barabensis). Gene 547, 273–279

42. Wang, Z., Jiang, S. F., Cao, J., Liu, K., Xu, S. H., Arfat, Y., Guo, Q. L., Chang, H., Goswami, N., Hinghofer-Szalkay, H., and Gao, Y. F. (2019) Novel findings on ultrastructural protection of skeletal muscle fibers during hibernation of Daurian ground squirrels: Mitochondria, nuclei, cytoskeleton, glycogen. J Cell Physiol

43. González, C. R., Muscarsel Isla, M. L., and Vitullo, A. D. (2018) The balance between apoptosis and autophagy regulates testis regression and recrudescence in the seasonal-breeding South American plains vizcacha, Lagostomus maximus. PloS one 13, e0191126–e0191126

44. Wang, Z., Jiang, S. F., Cao, J., Liu, K., Xu, S. H., Arfat, Y., Guo, Q. L., Chang, H., Goswami, N., Hinghofer-Szalkay, H., and Gao, Y. F. (2019) Novel findings on ultrastructural protection of skeletal muscle fibers during hibernation of Daurian ground squirrels: Mitochondria, nuclei, cytoskeleton, glycogen. J Cell Physiol 234, 13318–13331

45. Sato, T., Tachiwana, T., Takata, K., Tay, T. W., Ishii, M., Nakamura, R., Kimura, S., Kanai, Y., Kurohmaru, M., and Hayashi, Y. (2005) Testicular dynamics in Syrian hamsters exposed to both short photoperiod and low ambient temperature. Anatomia Histologia Embryologia 34, 220–224

46. Ushakova, M. V., Kropotkina, M. V., Feoktistova, N. Y., and Surov, A. V. (2012) Daily Torpor in Hamsters (Rodentia, Cricetinae). Russ. J. Ecol. 43, 62–66

